# Cellular labeling of endogenous virus replication (CLEVR) reveals de novo insertions of the gypsy endogenous retrovirus in cell culture and in both neurons and glial cells of aging fruit flies

**DOI:** 10.1101/445221

**Authors:** Yung-Heng Chang, Richard M. Keegan, Lisa Prazak, Josh Dubnau

**Affiliations:** Department of Anesthesiology, Stony Brook School of Medicine, NY 11794, USA; Program in Neuroscience, Department of Neurobiology and Behavior, Stony Brook University, NY 11794, USA; Biology, Farmingdale State College, NY 11735, USA

## Abstract

Evidence is rapidly mounting that transposable element expression and replication may impact biology more widely than previously thought. This includes potential effects on normal physiology of somatic tissues and dysfunctional impacts in diseases associated with aging such as cancer and neurodegeneration. Investigation of the biological impact of mobile elements in somatic cells will be greatly facilitated by use of donor elements that are engineered to report de novo events in vivo. In multicellular organisms, successful reporters of LINE element mobilization have been in use for some time, but similar strategies have not been developed to report Long Terminal Repeat (LTR) retrotransposons and endogenous retroviruses. We describe <u>C</u>ellular <u>L</u>abeling of <u>E</u>ndogenous <u>V</u>irus <u>R</u>eplication (CLEVR), which reports replication of the gypsy element in *Drosophila*. The gypsy-CLEVR reporter reveals gypsy replication both in cell culture and in individual neurons and glial cells of the aging adult fly. We also demonstrate that the gypsy-CLEVR replication rate is increased when the short interfering RNA silencing system is genetically disrupted. This CLEVR strategy makes use of universally conserved features of retroviruses and should be widely applicable to other LTR-retrotransposons, endogenous retroviruses and exogenous retroviruses.

## Introduction

Nearly 50% of the human DNA content, and equivalently vast fractions of most other animal and plant genomes, consist of sequences derived from transposable elements (TEs) [1]. TEs are selfish genetic elements whose primary adaptation is to copy themselves in germline tissue, thereby passing on de novo copies. Mutations generated by germline transposition are a significant source of genetic variability, with obvious impact on phenotypic diversity and on evolutionary adaptation [2, 3]. Although replication in germline is the means by which TEs generate inheritable de novo copies, it is increasingly clear that endogenous TEs also replicate in somatic tissues. The evidence for somatic transposition is particularly strong for Class 1 TEs, also known as retrotransposons (RTEs) [4-21]. RTEs replicate through an RNA intermediate by use of reverse transcription and insertion of de-novo cDNA copies. Somatic replication of RTEs has the potential for tremendous impact both on normal biological properties of tissues and on human health [2, 16-21].

Historically, most studies of the mechanisms of transposition have focused on events in the germline for two main reasons. First, replication in germline rather than somatic cells is the evolutionary adaptation that has permitted TEs to maintain their presence in the genome. So, from a conceptual point of view, it made great sense to study TE action there. The other reason why most research focused on germline events, however, is technical. Detection of de novo replication in germline is experimentally tractable because new insertions can be passed on to offspring where they are present in every cell. As a result, classical molecular approaches such as Southern blots as well as more modern genomic approaches are each feasible means to characterize new insertions. In contrast, detection of de-novo inserts in somatic tissues is far more challenging than in germline because each new TE insertion will be present in a single cell, or, at best, in one clonally related cell lineage if the insertions are followed by cell divisions. The discovery that LINE like RTEs are able to replicate during neural development in rodents [5-7, 14], as well as somatic cells during *Drosophila* embryonic development [15], however, provided the impetus to more closely examine TE replication in soma. It is increasingly clear that TEs also can be actively expressed and even mobile in somatic tissues such as brain during normal development [5-7, 9-14, 22-25], during aging in a variety of cell types and species [4, 26-31], in a variety of neurodegenerative diseases and in animal models of human neurodegeneration [32-41], and in cancer [17, 42-51].

Single cell whole genome sequencing approaches have provided the resolution to detect rare de novo insertions within individual somatic cells. As a whole, such studies have clearly supported the idea that some RTEs are capable of replication during brain development as well as in the context of cancer progression, although the rate of such de novo events and whether they have functional impacts remain controversial or unresolved [9, 10, 18, 23, 24, 43, 52-54]. But these approaches are limited in some important practical and conceptual ways. First, single cell sequencing is relatively expensive, and low throughput. In addition, the significant molecular biological and computational challenges that come with this approach have led to widely different estimates of mobilization frequency. But perhaps more importantly, detection of individual de novo replication events in single cells does not afford a platform to conduct structure/function manipulations of RTEs to dissect the biological mechanisms. Nor does it provide the opportunity to visualize or manipulate the cells in which the RTEs are replicating within a tissue.

A different approach to this problem is to engineer genetic reporters of RTE replication [4, 5, 55, 56]. Such reporters offer the advantage that they provide the means for structure-function studies to uncover cis-acting sites, investigation of impact of environmental or genetic perturbations, and to uncover the type and fate of the cells in which RTEs replicate within living tissues. Such reporters have been described for Ty LTR retrotransposons in yeast [55] and for L1 LINE elements in mammalian cell culture and in transgenic mice [5, 56]. Both of these reporters were constructed using the insertion of heterologous introns within the engineered RTE. The intron can be removed by splicing when the RTE passes through an RNA intermediate, thereby activating a reporter upon de novo integration of replicated cDNA copies into host DNA. The L1-GFP reporter mouse has revealed that de-novo mobilization events take place during mouse development [5] and provided the means to investigate the impact of behavioral and genetic perturbations [6, 7]. We previously described an indirect mobilization reporter for the gypsy-LTR retrotransposon/endogenous retrovirus (ERV). This so-called gypsy-trap made use of a known chromosomal hotspot for gypsy integration to turn off a Gal80 repressor upon insertion of a de novo retro-element into the hotspot cassette [4]. When present together with both a Gal4 activator and a Gal4 responsive GFP, this was sufficient to report integrations that disrupted the Gal80 cassette. This system has been successfully used to reveal de novo transposition during aging and in the context of neurodegeneration [4, 8, 26, 27, 36]. But there are several critical shortcomings of the gypsy-TRAP approach. First, it does not selectively report insertions of any single RTE. In fact, it can be activated by insertions from any gypsy-family member or indeed any RTE that shares that hotspot. Second it does not provide the means to manipulate the biology of the donor element. Third, because the gypsy-trap utilizes a repressor of Gal4 function, it precludes use of the powerful Gal4 system that is the primary tool for modern *Drosophila* genetics.

Although most of the work on somatic transposition has focused on LINE elements, an emerging literature has suggested the possibility of a far broader impact of RTEs within somatic tissues, with the possibility that both LINE-like and LTR RTEs and ERVs may mobilize [4-14, 25-51, 54]. But aside from the gypsy-Trap, whose utility is limited, no in vivo reporter system has yet been developed to monitor mobilization of donor LTR-retrotransposons or ERVs in any multi-cellular organism. Using the *Drosophila* ERV gypsy as a model, we have developed a novel reporter to reveal de novo mobilization both in cell culture and in somatic brain tissue. The gypsy-CLEVR reporter becomes hyper-active with disruption to the siRNA silencing system, which is known to help silence endogenous gypsy expression in somatic cells. Using this reporter system, we also confirm our previous report of age dependent increases in mobilization in neurons and demonstrate similar effects in glial cells. Because the CLEVR reporter system capitalizes on highly conserved mechanisms of replication, it likely will be generalizable to LTR-retrotransposons, endogenous retroviruses and even exogenous retroviruses in a variety of animal and plant species.

## Results

### Engineering of the *gypsy-CLEVR* reporter

To generate a reporter capable of tracing gypsy retrotransposition in specific cell lineages, we capitalized on conserved features of LTR-RTE/ERV replication (Fig 1A). Specifically, we took advantage of two universal features of retrovirus replication [57, 58]. First, because the promoter sequence in the 5’ end of the 5’LTR is not transcribed, the corresponding sequences at the 5’ end of the 3’LTR in the genomic RNA normally is used as a template to complete cDNA synthesis of the 5’ end of the transcript. Second, because the portion of the 3’ end of the 3’ LTR after the transcription termination signal is not transcribed, the corresponding portion of the 5’LTR is used to template replication of this region of the 3’LTR. As a result, each of the LTR sequences of the fully replicated cDNA are hybrids of the nucleotide sequence from the two LTR repeats. The CLEVR reporter takes advantage of this. We placed a promoter without a reporter at the appropriate (U5) position in the 3’ half of the 5’LTR and we positioned a reporter with no promoter into the appropriate (U3) location in the 5’ half of the 3’LTR. Thus, after full replication, the reinserted de novo element should have rearranged the promoter so that it is adjacent to the reporter at both the 5’ and 3’ LTRs.

**Fig 1.**
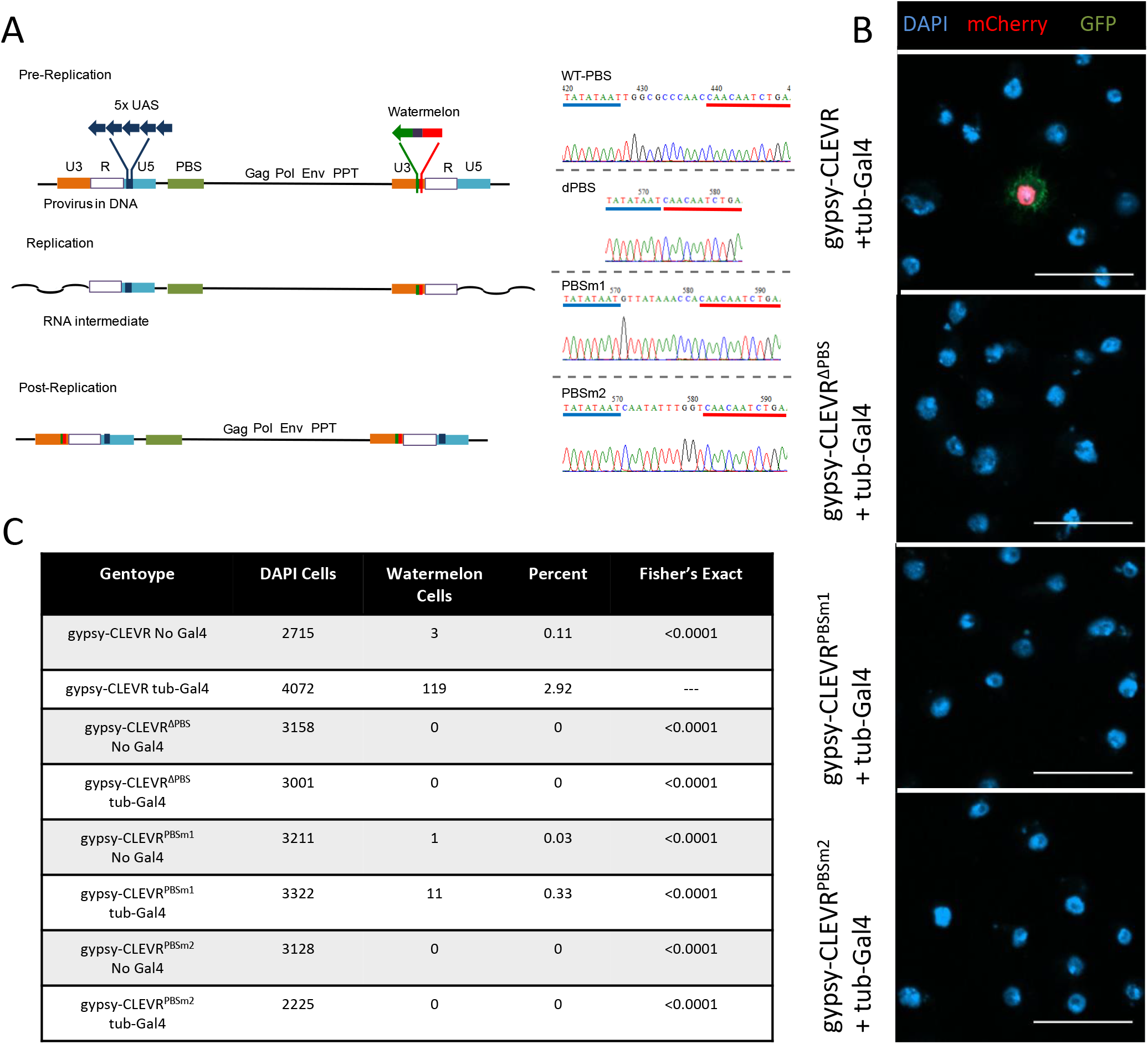
gypsy-CLEVR genetic structure and functional test in Drosophila S2 cells (A) Schematic showing the replication cycle of the gypsy-CLEVR reporter with the 5xUAS regulatory element and WM fluorescent reporter inserts in antisense orientation to gypsy. The wild type, deletion, and mutations of the PBS are shown with sequence confirmation. (B) Fluorescent images showing WM positive S2 cells for gypsy-CLEVR with tub-Gal4, but not with deleted (gypsy-CLEVR^ΔPBS^) or mutated (gypsy-CLEVR^PBSm1^ and gypsy-CLEVR^PBSm2^) primer binding sites. Scale bars = 20 μm. (C) Quantification of WM labeled cells with gypsy-CLEVR, gypsy-CLEVR^ΔPBS^, gypsy-CLEVR^PBSm1^, and gypsy-CLEVR^PBSm2^ with and without Gal4. Significance was determined by using a Fisher’s exact test comparing each genotype to gypsy-CLEVR with tub-Gal4.

In concept, CLEVR could use any promoter element that is small enough to fit in the 5’LTR without disruption of the ERV expression. In our case, we chose to use the Gal4 responsive 5XUAS enhancer, which is sufficient to drive strong expression under control of the Gal4 transcription factor. This promoter is compact in size and provides maximal functional utility because its use taps into a wealth of extant strains with Gal4 expression restricted to virtually any desired cell type or developmental stage. Importantly, the Gal4 responsive promoter faces to the left, opposite to the orientation of the gypsy promoter that lies in the LTR. Therefore gypsy expression falls under its own control. For a reporter, we wished to provide the means to clearly mark nuclei in order to count cells and identify the location of their soma within tissues. But we also wanted to be able to see cell morphology, for which it is helpful to highlight the plasma membrane. This is particularly important in brain tissue where glia and neurons can have extraordinarily complex morphologies. To accomplish both of these goals, we designed Watermelon (WM), a dual reporter module that encodes both a myristylated GFP that tethers to the membrane (myr-GFP-V5) and nuclear localized mCherry (H2B-mCherry-HA). The dual reporter expression relies on introduction of the porcine teschovirus-1 2A (P2A) self-cleaving peptide [59, 60] sequence (myr-GFP-V5-P2A-H2B-mCherry-HA, S1A Fig). When this dual reporter is expressed, it simultaneously marks the plasma membrane with GFP and cell nucleus with mCherry.

To test the functionality of the dual reporter, we created a UAS-WM construct, which we co-transfected with actin-Gal4 into *Drosophila* S2 cells. Indeed, this marked transfected cells with brilliant green membrane fluorescence surrounding bright red fluorescence in the nuclei (S1B Fig). We also generated UAS-WM transgenic flies, which we tested by crossing with several tissue specific Gal4 drivers. In the presence of *hedgehog (hh)-Gal4* (S1C Fig), the WM reporter marked posterior compartment cells [61] of wing discs during larval stage with mCherry in the nucleus and GFP on the membrane. We also tested this dual reporter in the *Drosophila* adult brain by expressing this transgene in subperineurial glia (SPG) under the control of the *moody-Gal4*. Here too, the nuclei were marked by strong mCherry fluorescence that co-localized with the glial nuclear Repo marker in SPG glial cells. The SPG glial membranes were painted with strong GFP, revealing the characteristic mesh-like architecture of this cell type (S1D Fig) [62]. These results demonstrate the functionality of the WM reporter to simultaneously mark cell nuclei (red) and membranes (green).

We next incorporated the dual label into the gypsy-CLEVR design (Fig 1A) and tested the reporter in S2 cells. When the gypsy-CLEVR construct was co-transfected along with tubulin-Gal4, strong WM fluorescent labeling was readily detected in approximately 3% of cells (Fig 1B and 1C). Within each labeled cell, the nuclei were painted with mCherry (which co-localized with DAPI) and were encircled by green fluorescence on the membrane. This is consistent with the idea that the gypsy-CLEVR reporter undergoes the rearrangement predicted during replication such that the promoter and reporter are placed adjacent to each other in the full-length cDNA. This rearrangement would then permit Gal4-mediated expression of the WM dual reporter. As predicted, WM labeling was not detected in the absence of Gal4 (Fig 1B and 1C). To rule out the possibility that the reporter can somehow be expressed in the absence of gypsy replication, we conducted a series of further experimental manipulations.

We first tested the impact of deleting (ΔPBS) or mutating (PBSm1 and PBSm2) the primer binding site (PBS), a cis-acting motif essential for retrovirus replication (Fig 1A). In contrast with the gypsy-CLEVER construct that contains the intact PBS, co-transfections of tubulin-Gal4 with the gypsy-CLEVR^ΔPBS^ did not yield any labeled cells (Fig 1B and 1C). Similarly, introduced mutations in the nucleotide sequence of the PBS greatly reduced (PBSm1; 0.33% of cells labeled) or eliminated (PBSm2; 0% of cells labeled) the reporter expression (Fig 1B and 1C).

### Molecular confirmation of *gypsy-CLEVR* retrotransposition *in vivo*

In order to test the efficacy and specificity of the gypsy-CLEVR reporter in vivo, we next introduced gypsy-CLEVR and its PBS modified variants (gypsy-CLEVRΔPBS, gypsy-CLEVR^PBSm1^ and gypsy-CLEVR^PBSm2^) into transgenic flies. We first sought to establish molecular confirmation that the donor RTE of the gypsy-CLEVR reporter was capable of replication, which should lead to the predicted rearrangement of the LTR sequences. The gypsy-CLEVR with an intact PBS and the variants with deletion or mutation of the essential PBS were tested in parallel. For each case, adult flies were collected directly after eclosing and were then aged for 7 days. Genomic DNA was extracted from these flies and nested PCR was performed to examine gypsy-CLEVR rearrangement. We did not detect visible signal in the 1st round of PCR (indicated region in Fig 2A). The DNA content in the target size region of the gel was extracted and used as template to perform a 2nd round of PCR. A PCR product of the predicted size was only detected in the intact PBS gypsy-CLEVR tissue, but not in non-transgenic control animals, and not in any of the three gypsy-CLEVR PBS modifying variants (Fig 2A). This PCR product was purified, sub-cloned and sequenced. The sequencing results from three independent clones confirmed the presence of the predicted rearrangement in the gypsy-CLEVR containing flies and the sequence of this clone matched that which is predicted to form after gypsy replication (S2A Fig). Together, these results provide strong molecular confirmation that the gypsy-CLEVR reporter is capable of replicating in vivo, and such replication is only detected at the PCR level in the presence of an intact PBS. We next tested whether the WM reporter could also be detected in these flies, and whether it too required an intact PBS.

**Fig 2.**
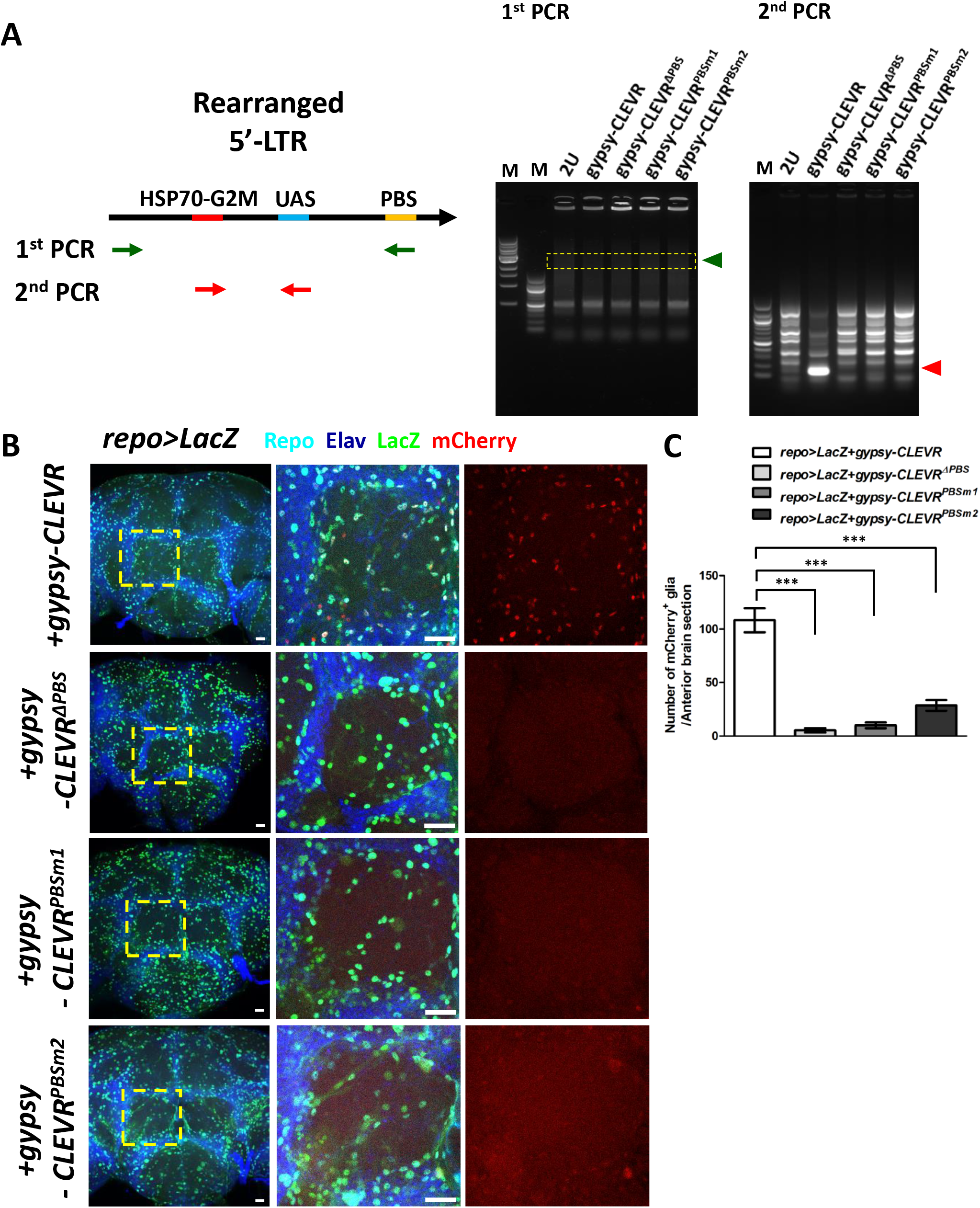
gypsy-CLEVR retrotransposition in glial cells vivo (A) Schematic illustration of LTR genetic structure after gypsy-CLEVR retrotransposition. Products of first and second rounds of nested PCR are shown from genomic DNA from 7 day old transgenic flies containing either wild-type gypsy-CLEVR, gypsy-CLEVR^ΔPBS^, gypsy-CLEVR^PBSm1^ or gypsy-CLEVR^PBSm2^. Results from first round of PCR (green primer set in schematic) do not detect a product. A second round of nested PCR was conducted from product isolated from highlighted gel region. Products of the correct size were detected (Red arrow) after this second round of PCR using nested primers. This product was only detected from flies with the intact gypsy-CLEVR transgene. PCR products were sequence verified (S2 Fig). (B) Gypsy-CLEVR replication was detected with confocal microscopy in adult brain glia by crossing flies containing the CLEVR constructs with a strain containing the pan-Glial Repo-Gal and a UAS-nuclearLacZ. CLEVR replication was detected using the mCherry nuclear signal (Red) as a reporter. Glial nuclei were independently labeled both with a repo antibody (Cyan) and with an antibody against beta-galactosidase (green). Neuronal nuclei were counter stained using the Elav antibody (Dark blue). Optical sections from anterior brain regions are shown (left panels) for 7 day old gypsy-CLEVR (wild-type, PBS deletion and mutations) transgenic flies. High magnification micrographs (from region indicated by yellow dash box) are shown to the right. Scale bar = 20 μm. (C) Quantification of the number of mCherry labeled glia per anterior brain section (Mean ± SEM) reveals that the mCherry labeling of glia requires an intact PBS (mCherry labeled glia for each group: gypsy-CLEVR: 108.3±11.2, gypsy-CLEVR ^Δ PBS^: 5.3±2.0, gypsy-CLEVR^PBSm1^: 10.0±2.7 and gypsy-CLEVR^PBSm2^: 28.7±5.0). Number of labeled glia is significantly higher with the wild type gypsy-CLEVR construct (*** p<0.001, unpaired t-test).

Because of our previous finding that *gypsy* expression and replication is increased with age [4] and because expression may be highest in glia [33], we first tested whether we could detect gypsy-CLEVR replication in glial cells of aged flies by crossing the reporter flies with the pan-glial driver, *repo-Gal4*. In 7-day old animals, we detect strong mCherry fluorescence in many glial nuclei and we detect GFP membrane fluorescence surrounding the nuclear cherry (Fig 2B and 2C; S3 Fig). By confocal imaging throughout the whole brain, we determined that most of the gypsy-CLEVR positive cells were located near the brain surface, especially in glia near the anterior regions (S4 Fig). We quantified the total number of WM labeled glia in anterior brain sections and found that, as was the case in cell culture, deletion or mutation of the PBS leads to a dramatic decrease in the number of labeled cells (Fig 2B and 2C), demonstrating that a cis acting motif that is essential for retrovirus replication also is required to activate the reporter. Although the replication of this construct is by design Gal4 independent, expression of the reporter is engineered to require Gal4. Indeed we rarely detected any gypsy-CLEVR positive cells in animals that did not contain the *repo-Gal4* driver, even when these flies were aged for 30 days (S5 Fig), a timepoint when endogenous gypsy is expressed at high levels [4, 33]. These rare labeled cells in flies that lack Gal4 likely derive from de-novo replication events that insert nearby active genomic loci, where the rearranged UAS-WM may be expressed by trapping nearby enhancers, thereby obviating the need for Gal4.

### Age dependent increase of gypsy retrotransposition in glial and neuronal cells

Our previous work using the gypsy-TRAP demonstrated an age-dependent increase in gypsy replication in neurons [4], and similar effects of age were subsequently documented with that reporter in other tissues [8, 26, 27, 36]. We therefore tested whether gypsy replication in glia was similarly age-dependent and could be detected using the gypsy-CLEVR reporter. We quantified the total number of gypsy-CLEVR positive glia in anterior brain sections and found that the number of labeled glia (6.5±0.9 per anterior brain) is extremely low in young animals, just 2 days after eclosion (Fig 3A and 3B). This number greatly increases (83.8±3.5 per anterior brain) by 5-7 days after eclosion (Fig 3A and 3B), consistent with the previous evidence that gypsy replication is age dependent in other cell types [4, 8, 26, 27, 36]. Moreover, this age-dependent increase in gypsy-CLEVR labeling could be blocked by introducing a Gal4 driven RNAi transgene against gypsy (UAS-gypsy-IR) [40] that we previously demonstrated was capable of interfering with the age-dependent increase in gypsy expression [33]. Addition of this UAS-gypsy-IR transgene was sufficient to block the age-dependent increase in gypsy-CLEVR labeled glia (34.5±5.4 per anterior brain) in 5-7 days old animals (Fig 3A and 3B). Thus, gypsy-CLEVR labeling requires gypsy expression as predicted.

**Fig 3.**
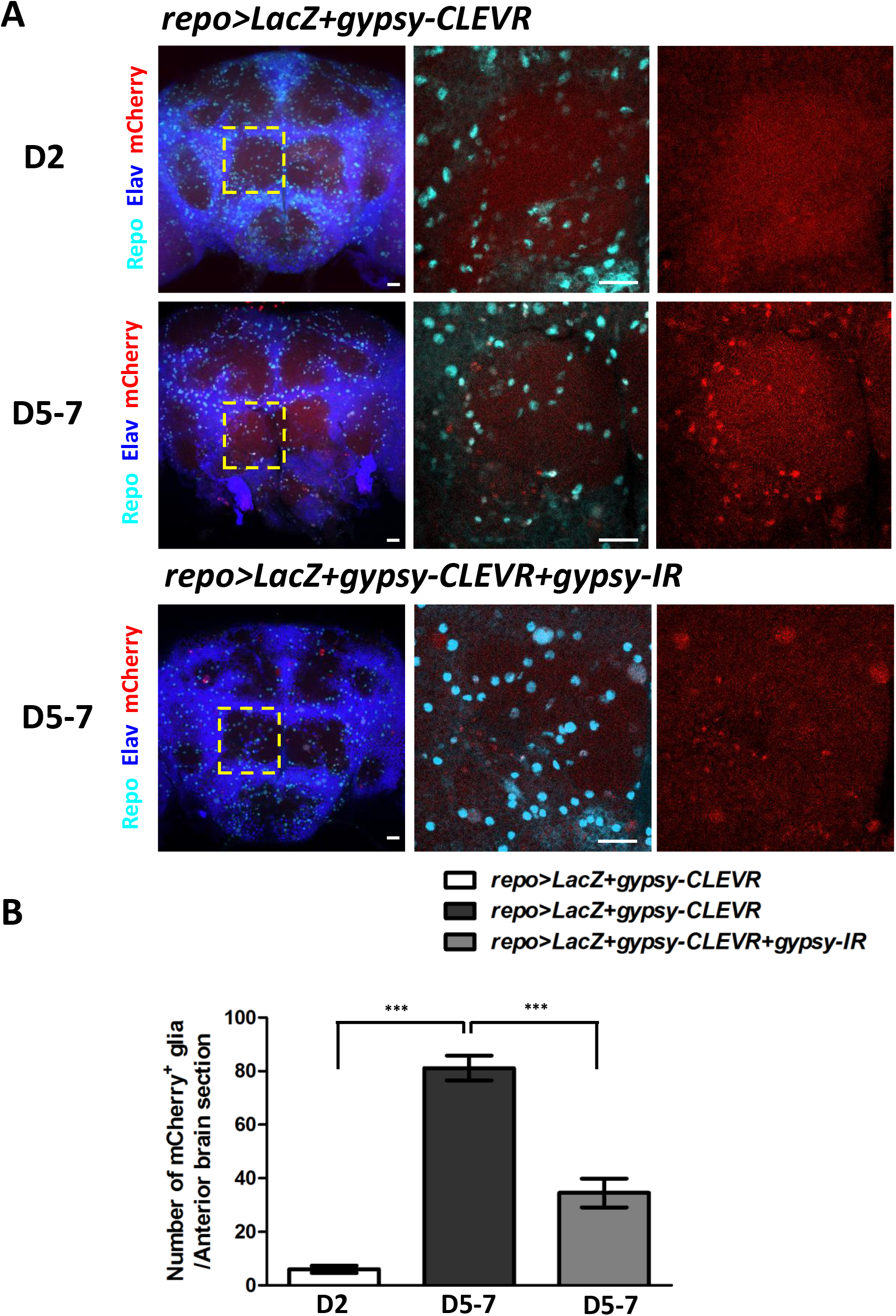
gypsy-CLEVR activity in glial cells is increased with age (A) Optical sections are shown from anterior brain regions from 2-day and 5-7 day old gypsy-CLEVR transgenic flies without and with a gypsy-IR transgene. Glial nuclei are independently labeled with antibodies against the glial Repo marker (cyan). Neuronal nuclei are labeled with antibodies against the Elav marker (blue) and gypsy-CLEVR replication is labeled with mCherry (red). High magnification images (Right panels) from within the regions marked with yellow dash box (left panels) are shown for each experimental group. Scale bar = 20 μm. (B) Numbers of mCherry labeled glial cells (Mean ± SEM) within anterior brain sections are shown for each experimental group (gypsy-CLEVR at 2 days: 6.5±0.9; gypsy-CLEVR at 5-7 days: 83.8±3.5; gypsy-CLEVR+gypsy-IR at 5-7 days: 34.5±5.4). (*** p<0.001, unpaired t-test).

We next tested the effects of age on gypsy-CLEVR labeling in neurons. Although our recent RNAseq experiments are consistent with the interpretation that gypsy levels are higher in glia than neurons [33], we previously demonstrated, using the gypsy-TRAP reagent, that gypsy is able to replicate in Kenyon cells (KCs), the intrinsic neurons of the mushroom body (MB), and that the number of de novo events increases with advancing age [4]. To test whether the gypsy-CLEVR reporter could also reveal age-dependent gypsy replication in these neurons, we used the *MB247-Gal4* line, which expresses in approximately 800 KC neurons per brain hemisphere [63]. Indeed, when we crossed *MB247-Gal4* to gypsy-CLEVR reporters (Fig 4), we detect rare de novo events in 2-day young animals (Fig 4A and 4G) and the population of gypsy-CLEVR expressing cells was significantly increased in 30-day old animals (Fig 4D and 4G). This result is consistent with our previous findings using the gypsy-TRAP reporter [4]. Importantly, the gypsy-CLEVR reporter is more sensitive (e.g. we detect events in younger animals) and more versatile in its applications. Together with the findings described above, and previously, these results demonstrate that gypsy retrotransposon activity is increased during aging in *Drosophila* adult brains, both in glia and neurons, and our gypsy-CLEVR reporter is capable to sensitively reveal these de novo retrotransposition events in vivo.

**Fig 4.**
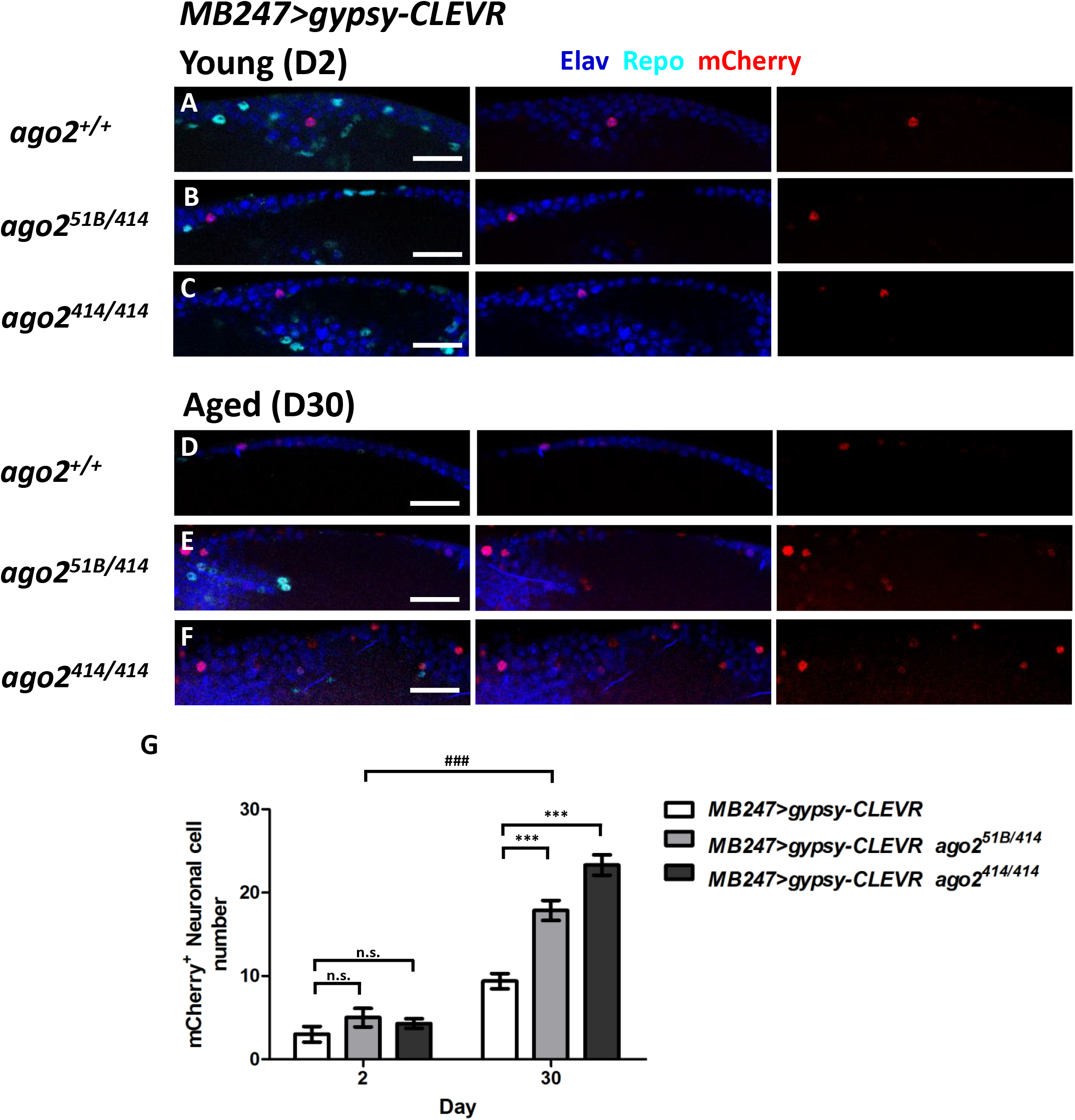
gypsy-CLEVR replication within KC neurons of the MB is increased with age, and with mutations in dAgo2. gypsy-CLEVR reporter was tested in flies that contain the *MB247-Gal4* which expresses in approximately 800 KC neurons of the Drosophila MB. Replication of the gypsy-CLEVR reporter was revealed by mCherry fluorescence (red). Neuronal nuclei were independently labeled with antibodies against the Elav marker (Blue) and glial nuclei were labeled with antibodies against the Repo marker (Cyan). Numbers of neurons labeled by gypsy-CLEVR reporter were significantly higher in brains from 30 day old vs 2 day old animals (A, D, G). Mutations in dAgo2 also significantly increased the number of mCherry labeled nuclei in 30 day old animals (D vs E and F) but not in 2 day old animals (A vs B and C). Scale bar = 20 μm. (G) Quantification of effects of age and genotype reveals a significant effect of age, and of mutations in dAgo2. (Mean ± SEM shown in bar graphs) Gypsy-CLEVR labeled cells for 2 day old animals: wild-type: 3.8±0.6, *ago2^51B/414^*: 5.0±1.1, *ago2^414/414^*: 4.3±0.6 and for 30 day old animals, wild-type: 9.4±0.9, *ago2^51B/414^*: 17.9±1.2, *ago2^414/414^*: 23.3±1.2). n.s., not statistically significant (unpaired t-test), *** p<0.001 (unpaired t-test), ### p<0.001 (two-way ANOVA).

### Argonaute-2 keeps gypsy replication in check in aging neurons

A previous literature has established that endogenous short interfering RNAs (siRNAs) loaded onto *Drosophila* argonaute-2 (dAgo2) plays a key role in transposon silencing in somatic cells [64-67], including brain [4]. Mutations in dAgo2 lead to increased expression of a suite of transposable elements in various somatic tissues [64-67] and leads to a precocious expression in brains of young animals of several RTEs including gypsy [4]. Our previous findings on the impact of dAgo2 on brain aging were consistent with the hypothesis that elevated gypsy expression could lead to an increased rate of mobilization, but we were not able to test this directly with the gypsy-Trap due to the practicalities of moving that multi-component system into homozygous dAgo2 mutant animals. We took advantage of the simplicity and sensitivity of the gypsy-CLEVR system to test whether mutations in dAgo2 impacted gypsy replication rate per se, rather than just expression.

We used the same *MB247-Gal4* driver to trace gypsy-CLEVR replication in MB KCs in *ago2* mutant animals. We find that in young flies at day 2, there is no significant increase in numbers of gypsy-CELVR positive Kenyon cell neurons in dAgo2 mutant (*ago2^51B/414^* and *ago2^414/414^*) vs. wild type (Fig 4A-4C and 4G) animals. But in brains from 30-day old animals, we detected a significant increase in numbers of gypsy-CLEVR positive KC neurons in both the *ago2^51B/414^* and the *ago2^414/414^* mutant genotypes compared to wild type controls (Fig 4D-4G). These data support the conclusion that the increase in gypsy expression levels in dAgo2 mutants leads to an increased rate of retrotransposition during aging in neurons.

## Discussion

The CLEVR reporter system relies on universally conserved features of retroviral replication to activate genetic reporter expression after successful mobilization of the donor element. We demonstrate that the gypsy-CLEVR construct reports gypsy replication events both in cell culture, and in several cell types in the adult *Drosophila* CNS. We present five separate lines of evidence that the activation of this reporter is caused by gypsy replication. First, we find that three different mutations to the PBS each are sufficient to disrupt reporter activation both in cell culture and in vivo. The PBS is a universally conserved and essential cis-acting site that is required to mediate priming of the first strand cDNA synthesis by tRNA (tRNA-Lys in the case of gypsy). This provides strong evidence that the activation of the dual reporter cannot occur in the absence of replication by some artifactual cause such as recombination. Second, we were able to verify with PCR and sequencing that the predicted replication-dependent rearrangement to place the UAS enhancer adjacent to the WM reporter actually takes place in vivo, and is not detectable when the PBS is disrupted. Third, using an RNAi transgene that targets gypsy sequences, we demonstrated that activation of the reporter requires expression of gypsy. Fourth, we found that the activation of the gypsy-CLEVR reporter is age dependent in neurons and in glial cells. This dovetails with a previous literature using the gypsy-TRAP reporter in which it was observed that gypsy de novo insertions accumulate with age in neurons [4, 36], adipose tissue [26, 27] and intestinal stem cells [8]. We further show evidence that the number of glial cells labeled by the gypsy-CLEVR reporter also increases with age. Fifth, we demonstrate using dAgo2 mutants that the gypsy-CLEVR reporter replication is inhibited by the siRNA surveillance system that normally stifles expression of endogenous gypsy elements. Taken together, these data strongly support the conclusion that CLEVR is both a sensitive and specific tool to reveal ERV replication in vivo.

The gypsy-CLEVR reporter offers significant advantages over our previously reported gypsy-TRAP. The TRAP reporter relied on using a DNA fragment containing a hot spot for gypsy family integration events, tethered to a Gal80 repressor in such a way that de novo integrations into that cassette would be likely to disrupt the Gal80 expression. In the presence of a Gal4 and a UAS-reporter, it was possible to reveal de novo events [4, 8, 26, 27, 36]. While this earlier reporter has utility, the CLEVR reporter offers a number of key improvements. First, the gypsy-TRAP does not distinguish which element has been inserted. In principle, any gypsy family member that shares a preference for the same hotspot can contribute labeled cells. Second, the gypsy-TRAP only captures the fraction of events that disrupt the particular cassette, and what fraction of the events are revealed is impossible to decipher. Third, the gypsy CLEVR system is far less cumbersome technically because it does not rely on three separate components like the gypsy-TRAP. Fourth, the gypsy-TRAP precludes the use of the Gal4 system to separately manipulate other pathways- e.g. we were able to use UAS-RNAi against gypsy to demonstrate specificity, which would not be possible with the gypsy-TRAP. Fifth, the CLEVR system affords the ability to conduct structure function studies such as manipulations of the PBS used here.

These features of the new reporter have already permitted us to investigate new aspects of gypsy biology. For example, we were able to demonstrate that dAgo2 not only represses gypsy expression in somatic cells as previously shown [4, 64-67], but that this has functionally relevant impact on the ability of gypsy to mobilize. This would have been difficult to test with the gypsy-TRAP because it would have necessitated combining 5 separate genetic components into one animal.

The CLEVR design in principle also should provide modularity needed to flexibly interrogate a diverse suite of research questions. In our case, we chose to use the UAS enhancer in the 5’LTR because it provided us the means to query gypsy donor mobilization in different cell types and under different conditions. For example, we were able to separately query glial cells or KC neurons by selecting different Gal4 drivers. In principle, however, any relatively small promoter sequence could be used. For example, if we wanted to use the Gal4 system to separately manipulate a genetic pathway of interest and then query the mobilization of gypsy within MB Kenyon cells, we could introduce the 247 base pair fragment of the dMef gene, which is sufficient to drive Gal4 in KCs (and is the basis of the *MB247-Gal4* line that we used here). CLEVR is also flexible with respect to the reporter in the 3’LTR. Here, we used a membrane GFP and a nuclear mCherry separated by the P2A sequence. It would be trivial to substitute reporters with different functional effects. For example, one could substitute GCaMP to image calcium or Channelrhodopsins for optogenetic manipulation to investigate functional correlates of gypsy mobilization in neurons. Finally, the CLEVR reporter relies on conserved features that are shared across all LTR-RTEs ERVs and exogenous retroviruses. Given the growing interest in the impact of ERVs on both normal and dysfunctional aspects of biology, the ability to follow insertion of proviruses within tissues should have palpable impact.

## Materials and Methods

### Constructs

To generate UAS-myr-GFP-V5-P2A-H2B-mCherry-HA (referred to as watermelon based on colors of fluorescence and abbreviated as WM), the myr-GFP and H2B-mCherry were separately amplified from pJFRC12-10xUAS-IVS-myr::GFP [68] and pEV-12xCSL-H2B-mCherry [69] vectors, and V5 and HA tags were separately added to C-terminals of myr-GFP and H2B-mCherry by polymerase chain reaction (PCR). To synthesize the final UAS-WM, the myr-GFP-V5 and H2B-mCherry-HA products were linked with P2A sequence [59, 60] and then inserted into the MCS of pUAST with NotI and XhoI digestion (see S1A Fig for full WM sequences). The gypsy backbone was obtained as a gift from V. Salenko [70]. In order to generate gypsy-CLEVR reporter, gypsy backbone that was placed in the pCaSpeR5 transforming vector [71], and the 5xUAS regulatory element from pUAST plasmid was inserted into the BglII site of U5 domain within gypsy 5’-LTR with antisense orientation relative to gypsy backbone. To synthesize the final gypsy-CLEVR reporter, the hsp70 promoter and the SV40 polyA tail from pUAST were separately fused upstream and downstream of WM reporter, and this final chimera was also placed in antisense orientation to gypsy transcript into XhoI site of U3 domain within 3’-LTR (see Fig 1 for gypsy-CLEVR structure in detail).

### Transgenic flies

All Gal4-drivers, UAS transgenes and mutants were backcrossed to our laboratory wild type strain, Canton-S derivative w^1118^ (*isoCJ1*) at least five generations. *MB247-Gal4, repo-Gal4, UAS-gypsy-IR, ago2^414^* and *ago2^51B^* [4, 33], *hh-Gal4* [61] and *moody-Gal4* [62] were gifts from published resources as indication. The *UAS-nls-lacZ* was obtained from Bloomington Drosophila Stock Center. Transgenic flies of *UAS-WM* and *gypsy-CLEVR* were generated by BestGene (CA, USA).

### Immunostaining of S2 cells and fly brains

*Drosophila* S2 cells (R69007, Thermo Fisher Scientific) were cultured in *Schneider’s Drosophila Media* (Thermo Fisher Scientific) supplemented with 10% Fetal Bovine Serum (Thermo Fisher Scientific) and Penicillin-Streptomycin-Glutamine (Thermo Fisher Scientific), in 75cm^2^ flasks. The actin-Gal4 [72] and tubulin-Gal4 [73] plasmids were used as previous publications. Cells were transfected with 1ug of each plasmid DNA with the Effectence transfection kit (Qiagen). After 48 hours transfection, cells were fixed in 4% Paraformaldehyde and mounted on coverslips coated in 0.5mg/ml Concanavalin A and ProLong Diamond Antifade Mountant with DAPI (Thermo Fisher Scientific).

Fly adult brains and 3rd instar wing discs were dissected, fixed and immunostained as previously described [4, 72]. The LacZ primary antibody was used at 1:500 dilution (A-11132, Thermo Fisher Scientific). Repo (8D12) and Elav (7E8A10) co-staining were performed using a 1:10 dilution (Developmental Studies Hybridoma Bank). DyLight 405, Alexa Fluor 488 and Alexa Fluor 647 conjugated secondary antibodies were used at 1:100 dilution to against Repo and Elav antibodies and at 1:500 to LacZ (Jackson ImmunoResearch). All S2 cell and fly brain images were acquired on a Zeiss LSM 800 confocal microscope and processed in Zeiss ZEN software package.

### Nested PCR and Sequencing

Genomic DNA was extracted from 20 flies of wild-type, gypsy-CLEVR and gypsy-CLEVR PBS modifying variants at day 7 by PureLink Genomic DNA Kit (Thermo Fisher Scientific). The extracted genomic DNA was followed by two rounds of standard PCR in the nested fashion. Primer 1 (5’-ACAATGTATTGCTTCGTAGC-3’) and primer 2 (5’-AGATTGTTGGTTGGGCGCCA-3’) were used in the first round PCR and the extract from first round PCR was amplified by primer 3 (5’-AAACTTAGTTTTCAATATTG-3’) and primer 4 (5’-ATCTGCTAGAGTCTCCGCTC-3’). The predicted size of PCR product from second round of PCR was extracted from gel and cloned by TOPO-TA cloning kit (Thermo Fisher Scientific). Sequencing was performed by DNA Sequencing Facility at Stony Brook School of Medicine.

### Statistical analysis

Cell culture data was analyzed using a Fisher’s Exact test variant of the Chi^2^ analysis in order to obtain a P value for significance. The statistical data from aging fly and genetic manipulation was analyzed by GraphPad Prism software. The unpaired t test was used to compare the different groups within same time point and two-way ANOVA was used to analyze the difference between groups with aging effect and genetic manipulation.

## Acknowledgments

We thank Prof. Jin Jiang (University of Texas Southwestern Medical Center) for the kind gift of *hh-Gal4*. We acknowledge Prof. Ulrike Gaul (Ludwig-Maximilians-University Munich) for sharing *moody-Gal4*. We thank Prof. Greg Beitel (Northwestern University) for sharing pCaSpeR5 vector, Prof. Paola Bellosta (University of Trento) for tubulin-Gal4 plasmid, Prof. Michael B. Elowitz (California Institute of Technology) for pEV-12xCSL-H2B-mCherry plasmid and Prof. Gerald M. Rubin (Janelia Research Campus) for pJFRC12-10xUAS-IVS-myr::GFP vector. We thank V. Salenko for the gypsy backbone vector. This work was supported by grants to J.D. from NINDS (R01NS091748) and the NIA (RF1AG057338).

## Supporting information

S1 Fig. Sequence information and functional test of WM transgene in S2 cell culture and different developmental stages of different tissues

(A) Sequence of WM (myr-GFP-V5-P2A-H2B-mCherry-HA). Myr-GFP sequence was shown in green with V5 tag in dark green. P2A sequence is shaded with yellow. H2B-mCherry shaded red with HA tag in pink. (B) *Drosophila* S2 cells were co-transfected with actin-Gal4 and UAS-WM (myr-GFP in green, H2B-mCherry in red and DAPI in blue). Scar bar = 10 μm. (C) Wing discs were dissected from flies with hh-Gal4 driving UAS-WM (*hh>WM*) at 3rd instar stage (myr-GFP in green, H2B-mCherry in red and DAPI in blue). Posterior cells expressing WM within yellow dashed box is shown in high magnification. Scar bar = 20 μm. (D) Moody Gal4 driving UAS-WM in SPG (*moody>WM*). Brains were dissected from adult flies and stained with glial marker Repo (magenta). Scar bar = 20 μm.

S2 Fig. Sequence confirmation of gypsy-CLEVR retrotransposition

Sequencing comparison of nested PCR products from 3 different batches of aged gypsy-CLEVR flies with comparison to predicted sequence of gypsy-CLEVR rearrangement after retrotransposition.

S3 Fig. The dual reporter labeling of nuclei and membrane can reveal cell morphology in vivo of cells in which gypsy-CLEVR replication has occurred.

The gypsy-CLEVR was separately crossed with *repo-Gal4* or *MB247-Gal4*. Adult brains from these crosses were dissected and labeled with glial marker (Repo in cyan), and both gypsy-CLEVR reporters, GFP (green, membrane) and mCherry (red, nuclei). Scale bar = 10 μm.

S4 Fig. Distribution of gypsy-CLEVR label in glial cells throughout the adult fly brain

Optical sections of 7 days old adult fly are shown from anterior, central and posterior regions. Glial nuclei are labeled with the pan glial marker Repo (cyan), neuronal marker, Elav (blue). Glial nuclei are independently labeled with a UAS-nuclearLacZ (green). gypsy-CLEVR reporter replication is revealed with nuclear mCherry (red). Highest levels of gypsy CLEVR replication are seen in anterior sections. Scale bar = 20 μm.

S5 Fig. Gal4 is required to activate WM reporter of gypsy-CLEVR

Gypsy-CLEVR transgenic flies that do not contain any Gal4 line were aged till 30 days, at which time high levels of gypsy expression and replication have taken place. Anterior brain sections of these aged gypsy-CLEVR adult contain few if any mCherry labeled nuclei (red). Glial nuclei were counter-stained with the Repo marker (cyan), neuronal marker Elav (blue). Scale bar = 20 μm.

**Author contributions:**
Conceptualization: Josh Dubnau
Data curation: Yung-Heng Chang, Richard M. Keegan
Formal analysis: Yung-Heng Chang, Richard M. Keegan
Funding acquisition: Josh Dubnau
Investigation: Yung-Heng Chang, Richard M. Keegan, Josh Dubnau
Methodology: Yung-Heng Chang, Richard M. Keegan, Josh Dubnau
Project administration: Yung-Heng Chang, Josh Dubnau
Resources: Yung-Heng Chang, Lisa Prazak
Software: Yung-Heng Chang, Richard M. Keegan
Supervision: Josh Dubnau
Validation: Yung-Heng Chang, Richard M. Keegan
Visualization: Yung-Heng Chang, Richard M. Keegan, Josh Dubnau
Writing – Original draft preparation: Yung-Heng Chang, Richard M. Keegan, Josh Dubnau
Writing – Review and editing:

